# An *Arabidopsis* receptor-like kinase mediates competitive plant-plant interactions

**DOI:** 10.1101/2025.08.10.667595

**Authors:** Cyril Libourel, Marie Invernizzi, Fabrice Roux, Mathieu Hanemian, Dominique Roby

**Affiliations:** Laboratoire des Interactions Plantes-Microbes Environnement (LIPME), INRAE, CNRS, Université de Toulouse, 31326 Castanet-Tolosan, France; Physiologie, Pathologie, et Génétique Végétales (PPGV), EI PURPAN, Université de Toulouse, Toulouse, France

**Keywords:** *Arabidopsis thaliana*, plant-plant interactions, receptor-like kinase, competitive response, natural variation

## Abstract

While competition among plant species is recognized as a major factor affecting crop yield and plant community dynamics, the genetic and molecular mechanisms underlying natural variation of such biotic interactions remain poorly characterized. Here, we report the cloning of a Quantitative Trait Locus previously detected in a Genome-Wide Association Study investigating the competitive response of *Arabidopsis thaliana* to the presence of the annual bluegrass weed species *Poa annua*. Using mutant and complementation strategies, we identified *ESCAPE 1* (*ESC1*) as the gene responsible for the natural variation of an escape strategy of *A. thaliana* in response to the presence of *P. annua*. *ESC1* encodes a proline-rich, extensin-like receptor kinase, also known as PERK13. An RNA-seq experiment revealed that PERK13 functions through different pathways in leaves and roots involving genes associated with responses to biotic and abiotic stresses. Using these RNA-seq together with yeast two-hybrid (Y2H) data, protein-protein interaction network reconstruction revealed two distinct decentralized protein networks in leaves and roots. These findings support the notion of an active response mechanism involved in neighbor detection. The functional validation of *ESC1* underlying natural variation in response to competition opens new avenues for a better understanding of the molecular dialogue involved in plant-plant interactions.

**Highlight:** In this study, we identify a receptor-like kinase enabling *Arabidopsis thaliana* to detect neighboring species through the activation of specific genetic pathways.

## Introduction

In natural environments, the structure, diversity, and dynamics of plant communities are largely shaped by competitive interactions among species (Tilman, 1985; Goldberg and Barton, 1992; Martorell and Freckleton, 2014). Similarly, in the absence of pesticides, competitive interactions between weeds and crop species are the primary biotic factor limiting crop biomass and grain yield, accounting for approximately 36% of losses, compared to about 18% caused by animal pests and 16% by pathogens (Oerke, 2006; Neve *et al*., 2009). Despite the importance of interspecific competition in the functioning of natural plant communities and crop performance, our understanding of the genetic and molecular bases underlying natural variation in interspecific competition is limited compared to other types of biotic interactions such as plant-pathogen interactions (Roux and Bergelson, 2016; Subrahmaniam *et al*., 2018; Becker *et al*., 2023).

To our knowledge, only four traditional Quantitative Trait Loci (QTL) mapping studies (using F2 or Recombinant Inbred Lines mapping populations) and four Genome-Wide Association mapping Studies (GWAS), reported the genetic architecture underlying the competitive response of a focal species to the presence of neighboring species (Coleman et al. 2001; Moncada et al. 2001; Granberry et al. 2016; Asif et al. 2015; Baron et al. 2015; Frachon et al. 2017; Libourel et al. 2021; Walsche et al. 2025). Across these studies, the genetic architecture was consistently polygenic, ranging from the identification of a few medium-effect QTLs to dozens of small-effect QTLs (Subrahmaniam *et al*., 2018; Sato and Wuest, 2025). In addition, the genetic architecture was strongly influenced by the identity of neighboring species, the diversity of surrounding species, and the composition of the plant assemblage (Libourel *et al*., 2021). In the four GWAS, all considering *Arabidopsis thaliana* as the focal species (Baron *et al*., 2015; Frachon *et al*., 2017; Libourel *et al*., 2021; Walsche *et al*., 2025), the fine mapping of genomic regions associated with natural variation in the response to the presence of neighboring species revealed numerous candidate genes related to key plant functions, including cell wall modification, signaling, and transport. Interestingly, these categories were not the most highly representedin experiments conducted under artificial conditions simulating plant-plant interactions, such as shading (Subrahmaniam *et al*., 2018). Furthermore, QTL cloning of genes associated with natural variation in interspecific competition has yet to be conducted, leaving the molecular mechanisms underlying this aspect of plant biotic interactions largely unexplored.

In this study, we identified and conducted a functional analysis of a gene underlying a QTL involved in the response to competition in *A. thaliana*. We used the data of a previous GWAS involving 91 accessions of *A. thaliana* from the highly genetically polymorphic French local mapping population TOU-A, grown either in the absence or in the presence of the bluegrass *Poa annua* (**Figure 1A**; Libourel et al. 2021)*. P. annua* is a common weed in cultivated fields (Li *et al*., 2009) and one of the primary grasses co-occurring with *A. thaliana* in natural plant communities and permanent meadows (Frachon *et al*., 2017, 2019). We targeted a QTL that explains approximately ∼15% of the genetic variation in both plant height from the soil to the first flower on the main stem and the height-to-diameter (HD) ratio, defined as the ratio between plant height and rosette diameter (**Figure 1B**; Libourel et al. 2021). High and low HD ratios have been shown to correspond to escape and aggressive strategies of *A. thaliana* in response to competition, respectively (Baron *et al*., 2015). Moreover, natural variation in the HD ratio in *A. thaliana* is under selection in the TOU-A population, which inhabits a highly competitive environment (Frachon *et al*., 2017). We therefore aimed to clone the causal gene *ESC1* underlying the QTL associated with the natural variation in the HD ratio. Among the genes located in this region, we demonstrated that *ESC1* corresponds to *PERK13* and mediates the escape strategy of *A. thaliana* in response to the presence of *P. annua*. In addition, through transcriptomics, yeast two-hybrid screening, and network reconstruction, we demonstrated that *ESC1/PERK13*-mediated pathways involve genes associated with both biotic and abiotic stress responses. Our findings suggest that plant responses to competition extend beyond competition for resources and engage molecular mechanisms similar to those involved in plant immune responses.

**Figure 1.**
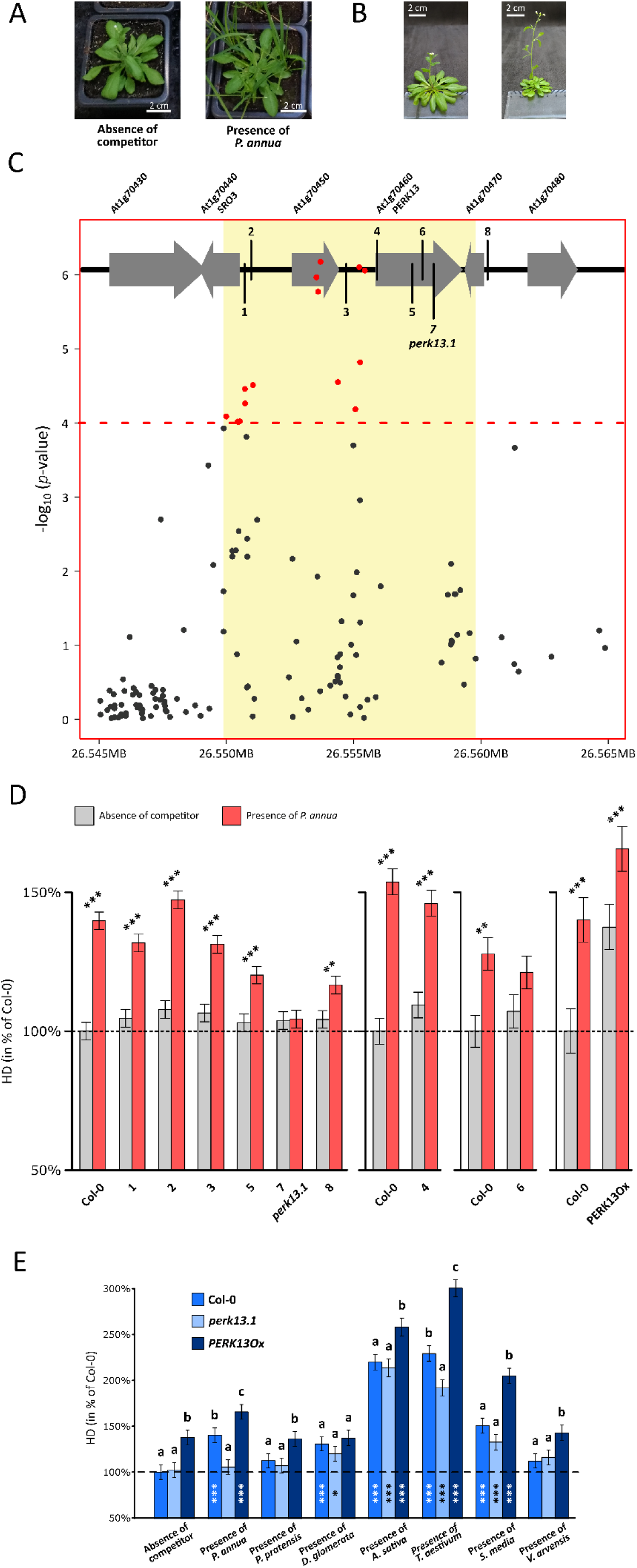
*ESC1/PERK13* contributes to the escape strategy of *A. thaliana* in response to *P. annua*. (A) Illustration of *A. thaliana* plants growing in the absence or the presence of *P. annua*. (B) Illustration of the natural variation of HD ratio illustrated by two accessions with contrasted phenotype, low HD ratio on the left and high HD ratio on the right. (C) Close-up view of the association peak identified for HD ratio in response to *P. annua* in Libourel et al., 2021, with a schematic representation of the genes located in the interval of the QTL identified. The numbered vertical bars represent the position of the selected mutant lines. The red dots indicate the most significant associated SNPs (-log_10_(*p*-value) > 4). The yellow rectangle indicates the confidence interval detected by a local score approach (Bonhomme et al., 2019). (D) Bar plots of the Least Square means (LSmeans) of the HD ratio from the tested *A. thaliana* genotypes in the absence (grey bars) or the presence (red bars) of *P. annua*. The values are expressed as a percentage of the HD ratio measured on Col-0in the absence of *P. annua*, with numbers corresponding to each mutant line. Phenotypes were obtained from at least three independent experiments. FDR corrected *p*-values: *0.05 > *P* > 0.01, **0.01 > *P* > 0.001, *** *P* < 0.001, absence of symbols: non-significant. (E) Bar plots illustrate the specificity of *ESC1*/*PERK13* towards other plant species besides *P. annua*. LSmeans of the HD ratio in the absence and presence of seven species (*P. annua*, *Poa pratensis*, *Dactylis glomerata*, *Avena sativa*, *Triticum aestivum*, *Stellaria media*, *Veronica arvensis*) expressed as a percentage of the HD ratio measured on Col-0 in absence of competitor. FDR corrected *p*-values for each genetic line between each treatment of interspecific competition and absence of competitor: *0.05 > *P* > 0.01, **0.01 > *P* > 0.001, *** *P* < 0.001, absence of symbols: non-significant. For each treatment, different letters indicate different groups according to the genetic lines after a FDR correction.

## Materials and Methods

### Plant materials

The T-DNA mutant lines of *A. thaliana* used in this study were ordered to the Nottingham *Arabidopsis* Stock Centre (NASC, http://Arabidopsis.info/BasicForm) and are in the Col-0 background (**Table S1**). One T-DNA mutant line is a GABI-Kat line (GK-345C10) whereas the remaining T-DNA mutants were identified in the SALK library (http://signal.salk.edu). The position of the T-DNA insertion was confirmed by polymerase chain reaction (PCR) using LP and RP primers designed using the online T-DNA Primer Design tools (http://signal.salk.edu/tdnaprimers.2.html) and the specific left border primer T-DNA insertion LBb1.3 (**Table S2**). Amplicons were sequenced using specific LP, RP, and LBb1.3 primers and assembled using Phred, Phrap, and Consed software. The results from sequencing were online BLAST using the web interface provided by NCBI (https://blast.ncbi.nlm.nih.gov/Blast.cgi) to identify the T-DNA insertional position in the genome. Seeds of the overexpressor line of *PERK13*, named *PERK13*Ox, were kindly provided by Prof. Hyung-Taeg Cho (Seoul National University, South Korea). This line, generated in the Col-0 background, contains the *PERK13* coding sequence under the control of the *EXPANSIN A7* promoter (Cho and Cosgrove, 2002) leading to an overexpression in root hairs. To reduce maternal effects, seeds of all these lines were produced under the same greenhouse conditions. For Y2H experiments, *Arabidopsis* Col-0 wild-type plants were sown on 100 squared petri dishes containing MS medium with the competitor *Poa annua* in a growth chamber at 21°C with a 16h photoperiod.

In this study, we also used the annual bluegrass *Poa annua* (Poaceae) as a neighboring species and six other species, namely the chickweed *Stellaria media* (Caryophyllaceae), the speedwell *Veronica arvensis* (Plantaginaceae), the Kentucky bluegrass *Poa pratensis* (Poaceae), the cat grass *Dactylis glomerata* (Poaceae), the oat *Avena sativa* (Poaceae), the common wheat *Triticum aestivum* (Poaceae) (**Table S3**). Seeds for the first four species were obtained from the Arbiotech Company (http://www.arbiotech.com). Seeds for *A. sativa* and *T. aestivum* were kindly provided by Dr. Etienne-Pascal Journet (AGIR, INRAE, Castanet-Tolosan, France).

### *PERK13* sequencing, plasmid constructions, and transgenic plant generation

The *PERK13* gene and its flanking regions (∼5.6kb) from the Col-0 accession and eight accessions of the local TOU-A mapping population (A1-115, A1-79, A6-104, A1-124, A6-61, A6-107, A1-120 and A6-27, **Figure 2A**), were sequenced after amplification with the RHS10_LR_Fwd and RHS10_LR_Rv primers using the PrimeSTAR® GXL DNA Polymerase (Takara) (**Table S2**). The sequencing was performed by Sanger technology using 23 primers (RHS10_LR_X) to cover the 5.6kb region (**Table S2 & Supplementary File 1**). Sequences were assembled using the Phred, Phrap, and Consed softwares. The results were online BLAST using the web interface provided by NCBI (https://blast.ncbi.nlm.nih.gov/Blast.cgi).

**Figure 2.**
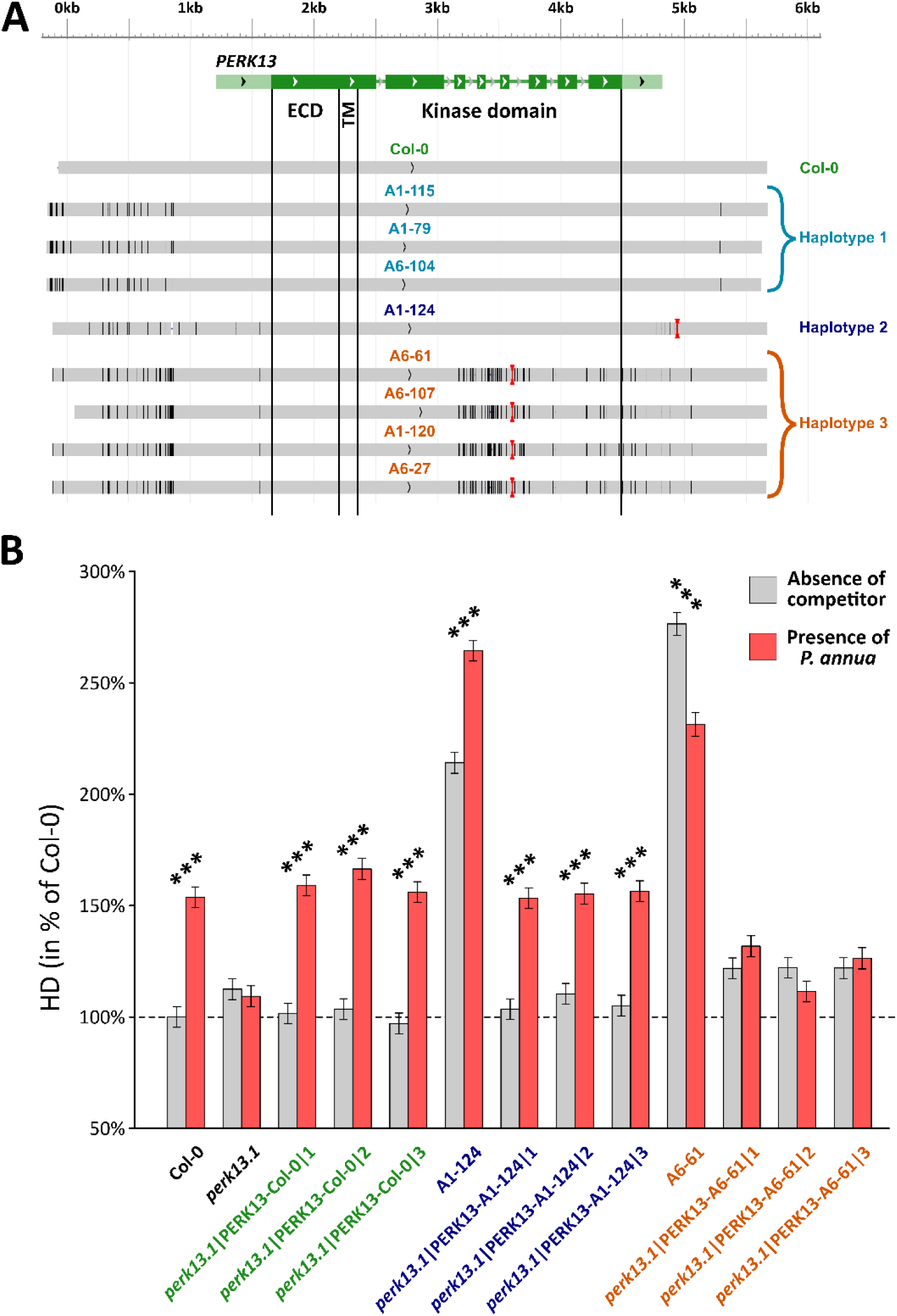
*ESC1/PERK13* genetic diversity controls the natural variation of *A. thaliana* response to the presence of *P. annua*. (A) Sequence diversity observed in a ∼5.6kb region centered on *PERK13* in Col-0 and eight accessions from the TOU-A population. Black vertical lines indicate mismatches, and white vertical lines indicate deletions. Insertions are represented by red hourglasses. (B) Barplots of the Least Square means of the HD ratio in the absence and presence of *P. annua* expressed as a percentage of Col-0 in the absence of *P. annua*. FDR corrected *p*-values between the absence and presence of *P. annua* for each genotype: *0.05 > *P* > 0.01, **0.01 > *P* > 0.001, *** *P* < 0.001, absence of symbols: non-significant.

To generate the constructs for complementation experiments, amplicons obtained for Col-0, A1-124, and A6-61 with the primers attB1_primer and attB2R_primer (**Table S2**) were cloned into the donor vector pDONR207, using multisite Gateway technology (Life Technologies). Subsequently, the respective constructs were cloned into the pEG301 vector and introduced in *Agrobacterium tumefaciens* (strain GV3101) by electroporation. Three-week-old *perk13.1* loss-of-function plants were transformed by floral dip (Clough and Bent, 1998). For each construct, at least three independent homozygous lines were selected for phenotyping and molecular characterization.

### Measurement of aboveground traits

#### Experimental design

Phenotyping experiments were replicated three times for each line used in this study (**Table S4**). For phenotyping of the T-DNA mutant lines and the complemented lines, we used for each replicate a split-plot design arranged as a randomized complete block design (RCBD) with two competition treatments (i.e. absence and presence of *P. annua*) nested within blocks (**Table S4**). We included Col-0 as a control in each experiment (**Table S4**). For testing the specificity of the competitive response mediated by *PERK13* towards other plant species than *P. annua*, we used a split-plot design arranged as a randomized complete block design (RCBD) with eight competition treatments (i.e. absence and presence of seven different species, nested within blocks (**Table S5**).

#### Growth conditions

The experiments were conducted in the same growth chamber of the Toulouse Plant-Microbe Phenotyping Platform (TPMP, https://eng-phenotoul.hub.inrae.fr/who-we-are/phenotoul/tpmp). Pots (7cm x 7cm x 6cm) were filled with damp standard culture soil (PROVEEN MOTTE 20, Soprimex). In presence of neighboring species, each *A. thaliana* plant was surrounded by three neighboring plants. Seeds for neighboring plants were evenly spaced, 2 cm away from the *A. thaliana* central position. Both species are sown the same day. During the experiments, plants were grown at 20 °C under artificial light to provide a 16-hr photoperiod and were bottom watered without supplemental nutrients. *A. thaliana* focal seedlings and *P. annua* seedlings were thinned to one or three per pot respectively, 6 to 12 days after seed sowing. The germination date of *A. thaliana* target seedlings was daily monitored.

#### Phenotypic traits

Two raw phenotypic traits were measured on each focal plant of *A. thaliana* at the time of their flowering, which was measured as the number of days between germination and flowering date. The first trait corresponds to the height from the soil to the first flower on the main stem (H1F expressed in mm). H1F is related to seed dispersal (Wender *et al*., 2005) and shade avoidance (Dorn *et al*., 2000) in *A. thaliana*. The second trait corresponds to the maximum diameter of the rosette, which was measured at the nearest millimeter (DIAM) (Weinig *et al*., 2006). This trait is a proxy of the growth of the rosette of the focal plant from germination to flowering. These traits allowed us to estimate the HD ratio as the height from the soil to the first flower on the rosette diameter (i.e. H1F/DIAM).

#### Statistical analysis

The following mixed model (PROC MIXED procedure, REML method, SAS 9.3, SAS Institute Inc) was used to explore the phenotypic differences among the different lines:

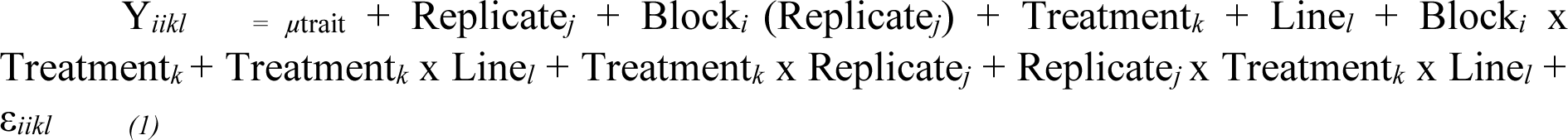

where ‘*Y*’ is the HD ratio scored on focal *A. thaliana* plants, ‘µ’ is the overall phenotypic mean; ‘Replicate’ accounts for differences among the temporal replicates; ‘Block’ accounts for environmental variation among experimental blocks within each replicate; ‘Treatment’ corresponds to the effect of the presence of a neighboring species (absence of competitor *vs* presence of *P. annua, V. arvensis, S. media, P. pratensis, D. glomerata, A. sativa,* and *T. aestivum*); ‘Line’ measures the effect of the different genetic lines; the interaction term ‘Treatment’ x Line’ accounts for variation among genetic lines for their reaction norms across the treatments. All factors were treated as fixed effects. For the calculation of *F*-values, terms were tested over their appropriate denominators. Given the split-plot design used in this study, the variance associated with ‘Block x Treatment’ was used as the error term for testing the ‘Block’ and ‘Treatment’ effects. Least Square means (LSmeans) of the HD ratio were obtained for each ‘Treatment x Line’ combination following the model described above (Supplementary_dataset 1).

### Y2H library construction and screening

*Saccharomyces cerevisiae* strains Y2H Gold and Y187 were grown at 30°C on YPDA medium and were used for bait and prey library cloning respectively. RNA extraction was performed from *A. thaliana* plants (Col-0) harvested at 4, 10 and 14 days (at least 3 plants/sample) after sowing and in presence of the plant competitor *P. annua* (3 independent experiments). RNA quality and integrity were checked using RNA 6000 Nano chip (Agilent, A_260_/A_280_ ∼2, A_260_/A_230_ = 2.0-2.2 and RIN>6). The library construction was performed from RNA samples from each timepoint, which were subsequently pooled in equimolar ratio and using the Make Your Own “Mate & Plate™” Library System (Takarabio, 2021). mRNA was purified using Invitrogen Dynabeads™ Oligo(dt)_25_ following the protocol supplied by the manufacturer. The cDNA mix was co-transformed with pGADT7-Rec plasmid in competent Y187 cells using the Yeastmaker Yeast Transformation System 2 (Takarabio, 2010).

For library screening, the PERK13 Kinase domain (KD) was used after PCR amplification (**Table S2**) for cloning into pGBG-GWY expression vector and transformation of *S. cerevisiae* Y2HGold strain (AUR1-C, ADE2, HIS3, and MEL1). Yeast transformants were selected on SD media lacking tryptophan, leucine and histidine (-TLH) and lacking also Adenine (-TLHA). Candidate interactors were identified by sequencing using specific pGAT-T7-Rec primers (**Table S2**). For one-by-one Y2H interactions, plasmids of the candidates were co-transformed with PERK13-KD in Y2H Gold yeasts on SD-LT media and then transferred on SD-TLH media.

### Gene expression analysis by RT-qPCR

Total RNA was extracted from 14-day-old plant roots grown under *in vitro* conditions, from eight to nine biological replicates per genotype and three independent experiments, with the NucleoSpin® RNA kit (Macherey Nagel). 500ng of total RNA was used for cDNA synthesis with the reverse transcriptase Transcriptor according to the manufacturer’s protocol (Roche E1372). Quantitative RT-PCR reactions were performed in 10µL using SYBR ® Green II master mix (Brilliant II SYBR Green QPCR Master Mix, Sigma-Aldrich). Gene expression was normalized using the housekeeping genes *ACT2* (AT3G18780) and *MON1* (AT2G28390) (Czechowski *et al*., 2005).

### Transcriptomic analysis

#### Sample preparation and total RNA extraction

Leaves and roots of *perk13.1*, PERK13Ox and Col-0 lines grown in the presence or absence of *Poa annua*, were sampled at 9, 15 and 21 days after sowing and flash frozen in liquid nitrogen. Three replicates of 3 pooled plants were sampled for each timepoint except for the 1st timepoint, where 12 plants were pooled. Three independent experiments were conducted, and total RNA extraction was performed using Nucleospin® RNA plus kit (Macherey-Nagel) and quality was checked using RNA 6000 Nano Kit (Agilent).

#### RNA sequencing and analysis

RNA sequencing was outsourced to Novogene (Cambridge, United Kingdom) to produce Illumina 150 bp paired-end Illumina using NovaSeq6000 sequencing platform. Raw paired-end reads were processed using the nf-core/rnaseq pipeline version 3.0 (doi:10.5281/zenodo.4323183). Pseudo-mapping strategy with salmon software (version 1.4.0) and reference genome annotation of *Arabidopsis thaliana* version Araport11 were used. Counts were retrieved at the gene level and normalized using the TMM method from EdgeR package (version 3.38.4). Differential gene expression analysis was performed using R version 4.2.0 (2022-04-22) and the EdgeR_3.40.2 (Robinson *et al*., 2010). DEG lists were generated by comparing expressions of *perk13.1* or PERK13Ox lines to Col-0, for each treatment and time point. From these lists, genes with an FDR < 0,1 were selected and joined to create a new DEGs list at 21 das (**Table S6**). The packages ggplot2 v.3.4.3, ggpubr v.0.6.0 and vegan 2.6.4 were used to generate the graphs (Wickham, 2016; Oksanen *et al*., 2022; Kassambara, 2023). Gene Ontology analysis was done using Classification SuperViewer Tool w/ Bootstrap (Provart & Zhu, 2003). All GO terms associated with selected DEGs can be found in **Table S6**.

#### Network reconstruction

BioGRID protein interaction dataset version 4.4.227 was used to recover interactors of the 503 deregulated genes found by transcriptomic analysis, and of the 14 candidate interactors of PERK13 identified by Yeast two-Hybrid screens. Cytoscape software v3.10.1 was used to plot protein-protein interactions, and cytoHubba application was used to calculate the network connectivity (Maere *et al*., 2005; Lin *et al*., 2008; Chin *et al*., 2014). Data used to build the networks are found in the **Table S7**.

## Results

### *ESC1* corresponds to *PERK13* and mediates the competitive response to *Poa annua*

To investigate the molecular mechanisms involved in the natural variation of *A. thaliana* competitive response to *P. annua* (**Figure 1A** and **1B**), we aimed at cloning the gene underlying a QTL explaining ∼15% of the genetic variation in the HD ratio (Libourel *et al*., 2021), hereafter called *ESCAPE1* (*ESC1*). A close-up revealed that this QTL, located at the end of chromosome 1, corresponds to a neat association peak spanning ∼20kb (**Figure 1C**). This short genomic region includes four genes, namely *AT1G70440* (*SIMILAR TO RCD ONE3*)*, AT1G70450*, *AT1G70460* (*PROLINE RICH, EXTENSIN-LIKE RECEPTOR KINASE 13* - *PERK13*, also named *ROOT HAIR SPECIFIC 10* - *RHS10*), and *AT1G70470*.

To identify the gene underlying the natural genetic variation in the HD ratio in response to *P. annua*, we measured the phenotypic response to the presence of *P. annua* of eight T-DNA mutant lines corresponding to the genes located within the 20kb region of the QTL (**Figure 1C** and **Table S1**). The wild-type Col-0 accession exhibited a significant increase (+40%) in the HD ratio in response to the presence of *P. annua* compared to the control condition (**Figure 1D**). Seven T-DNA mutants exhibited a response similar to Col-0, although the increase in HD ratio observed in mutant #*6* was not statistically significant (**Figures 1D, Table S8**). On the other hand, no increase in the HD ratio was observed for the loss-of-function mutant #7 (hereafter named *perk13.1*), suggesting that *PERK13* plays a major role in the competitive response to *P. annua*. In line, a genotype overexpressing *PERK13* in root hairs, *PERK13*Ox (Hwang *et al*., 2016), exhibited a constitutive higher HD ratio than Col-0 in the absence of *P. annua* as well as an increase in the HD ratio in response to the presence of *P. annua*.

A functional complementation of the *perk13.1* mutant with a ∼5.6kb genomic region including the Col-0 *PERK13* allele in three independent complemented lines (*perk13.1*|*PERK13*-Col-0) led to a complete restoration of the response to wild-type level (**Figures 2B**, **Table S9**). Together, these results demonstrate that *ESC1*/*PERK13* plays a significant role in the competitive response of *A. thaliana* to the presence of *P. annua* by mediating an above-ground escape strategy.

Then, we explored the specificity of *ESC1/PERK13* in the competitive response by growing Col-0, *perk13.1,* and *PERK13*Ox lines in the presence of *P. annua*, as well as four other grass species (including two wild species and two crops, **Table S3**), and two herb species commonly associated with *A. thaliana* in natural plant communities (**Figure 1E** and **Table S5)**. We observed similar phenotypic responses between Col-0 and *perk13.1* across most herb and grass species with an increase of HD ratio (Figure 1E and Table S5 and Supplementary_dataset1). However, in the presence of the common wheat *T. aestivum*, *perk13.1* displayed a lower HD ratio than Col-0, while *PERK13*Ox showed a higher HD ratio, therefore mirroring the response observed with *P. annua*. These results indicate that the role of *PERK13* in the competitive response of *A. thaliana* depends on the identity of the neighboring species.

### The competitive response to *P. annua* depends on *PERK13* sequence variability in natural accessions

To gain insights into a potential relationship between *ESC1/PERK13* natural sequence diversity and its role in response to *P. annua* presence, we selected eight TOU-A accessions used in the initial GWAS. We chose them according to their contrasting response to *P. annua* as well as their allele for the most associated SNP at the position 26,555,224 on chromosome 1, with four accessions chosen for each allele (**Figure 2A**). The sequencing of the eight TOU-A accessions and the re-sequencing of Col-0 for a ∼5.6kb region encompassing the promoter and the coding regions of *PERK13*, revealed 93 indels and 88 SNPs (**Figure 2A**). The majority of polymorphisms were located in the kinase domain (49%) and the promoter region (38%), while none were detected in the transmembrane or extracellular domains of *PERK13* (**Figure 2A**). Strikingly, the 17 polymorphisms located in the exons correspond to synonymous mutations. We identified three distinct haplotypes among the eight TOU-A accessions (**Figure 2A**). While Haplotype 1 and Haplotype 2 are closely related to Col-0, Haplotype 3 differs significantly from them, particularly in the kinase domain region, where the 88 identified polymorphisms are in complete linkage disequilibrium.

To test whether these highly divergent haplotypes mediate different competitive responses to the presence of *P. annua*, we complemented the mutant *perk13.1* with *PERK13*-Haplotype 2 and *PERK13*-Haplotype 3 (**Figures 2B**). We observed that the three lines complemented with *PERK13*-Haplotype2 showed a significant increase in the HD ratio in response to the presence of *P. annua*, similar to Col-0 or the lines complemented with the Col-0 haplotype of *PERK13* (**Figure 2B**). In contrast, the three lines complemented with *PERK13*-Haplotype3 exhibited similar HD ratios in both the absence and presence of *P. annua* (**Figures 2B**, **Table S9**. Together, these results demonstrate the existence of a functional haplotype (Haplotype 2) and a defective haplotype (Haplotype 3), thereby confirming the role of *PERK13/ESC-1* in the natural variation of the HD ratio in response to the presence of *Poa annua*.

### The *ESC1/PERK13*-dependent response to *P. annua* competition relies on stress-related genes

To investigate the molecular pathways involved in the escape strategy mediated by *ESC1/PERK13*, we conducted an RNAseq experiment on 9, 15, and 21 day-old *A. thaliana* roots or leaves of *perk13.1*, *PERK13*Ox, and Col-0 grown in the presence or absence of *P. annua*. We compared the transcriptional reprogramming in *perk13.1* or *PERK13*Ox to the transcriptional reprogramming in Col-0 at each developmental stage and in each compartment to identify differentially expressed genes (DEGs). In the presence of *P. annua*, we observed distinct transcriptomic changes between leaves and roots in the *perk13.1* mutant. For instance, in 21-day-old plants grown in the presence of *P. annua*, 462 DEGs were identified in the leaf compartment, while only 57 DEGs were identified in the root compartment. Notably, only 5 genes were identified as commonly deregulated in the two compartments in response to *P. annua* (**Figure 3A** and **3B**). Moreover, while the number of DEGs across development stages was relatively stable in the root compartment, regardless of the absence or presence of *P. annua*, a sharp increase in the number of DEGs was observed in the leaf compartment during the development of *A. thaliana*, especially in presence of *P. annua* (**Figure 3A**). Compared to the *perk13.1* mutant, we observed a higher number of DEGs for the overexpressing line in the two plant compartments whatever the absence or presence of *P. annua* (**Figures S1** and **S2**). Consequently, we focused our further analysis on the DEGs of the *perk13.1* mutant in 21-day-old plants, since most of the genes were deregulated at this developmental stage. In addition, we only kept the genes differentially expressed specifically in the presence of *P. annua*. This led to the final sets of 458 and 45 DEGs in the leaf and root compartments, respectively (**Figure 3B**).

**Figure 3.**
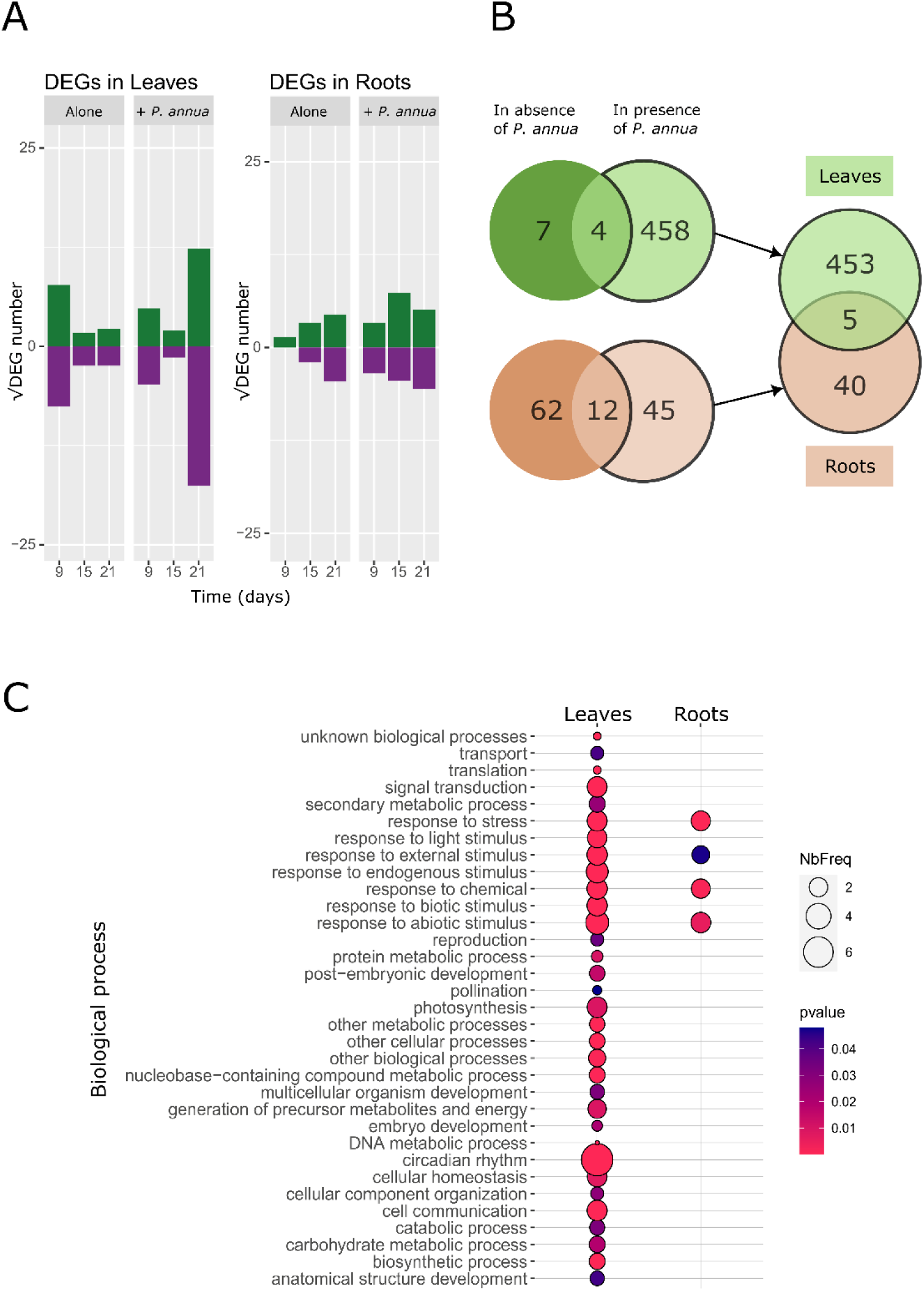
Transcriptional reprogramming in *perk13.1* in response to *P. annua*. (A) Upregulated genes (green) and downregulated genes (purple) found in the *perk13.1* mutant in comparison to Col-0 for each treatment, tissue, and timepoint. (B) Venn diagrams representing the DEGs found in 21-day-old plants in the absence (dark color) and the presence of *P. annua* (light color), for the leaf compartment (green) and the root compartment (orange). The parts of the diagram outlined in bold represent the DEGs specific to *P. annua* response and are used for DEGs comparison between the leaf and root compartments in (C). (C) Go terms enriched in leaves and roots of 21-day-old plants of the *perk13.1* mutant in the presence of *P. annua*. *P*-values < 0.05 were selected and the Normed Frequency (NbFreq) was calculated as follows: (Number_in_Class input/Number_Classified input)/(Number_in_Class reference/Number_Classified reference).

To understand more precisely the molecular pathways driven by *ESC1*/*PERK13* in response to *P. annua,* a GO term enrichment analysis was performed on these two lists of genes revealing a global reprogramming primarily centered on stress responses. In the leaf compartment, while DEGs were related to diverse biological processes responses such as circadian rhythm and cell communication, a clear enrichment was observed in many categories related to the response to either biotic or abiotic stimuli (**Figure 3C**). In the root compartment, a few functional categories also related to stress were enriched. Interestingly, in both compartments, several DEGs were associated with plant immune responses, including (i) perception related genes such as LRR protein kinases, (ii) signaling related genes including kinases, calcium and ROS related genes, (iii) transcription factors, and (iv) genes related to metabolism and hormone regulation (**Table S6**).

### ESC1/PERK13 is part of two decentralized protein networks

Our transcriptomic analysis allowed the identification of *ESC1/PERK13*-dependent genes that were deregulated in response to the presence of *P. annua* and that might be important for setting up an escape strategy. To gain a broader understanding of the complex response of *A. thaliana* to the presence of *P. annua* and identify key regulatory components, we reconstructed protein–protein interaction networks using the sets of DEGs identified in our transcriptomic analysis conducted on the leaf and root compartments. In addition, we performed a Yeast-Two-Hybrid (Y2H) screen using a library generated from *A. thaliana* plants grown in the presence of *P. annua*, enabling the identification of candidate proteins interacting with the ESC1/PERK13 kinase domain. Among the 15 candidate proteins identified (**Table S10**), four of them were identified at high frequency and correspond to two calcium-related proteins [CALMODULIN-BINDING RECEPTO-LIKE CYTOPLASMIC KINASE 2 (CRCK2) and CDPK-RELATED PROTEIN KINASE 4 (CRK4)], a VASCULAR PLANT ONE ZINC FINGER PROTEIN 1 (VOZ1), and LYSOPHOSPHOLIPASE 2 (LysoPL2). Reconstruction of our interaction networks was performed using these candidate proteins, along with the DEGs, and by looking for known interactors of the proteins encoded by the DEGs and of the ESC1/PERK13 putative interactors identified in this study. As RP1 (At1g43170) is a ribosomal subunit, a family reported as a classic false-positive interactor in Y2H assays (Van Criekinge and Beyaert, 1999), it was excluded from further analysis.

The leaf protein-protein interaction (PPI) network consists of 3032 interactions between 1502 proteins (nodes) (**Figure 4**, **Table S7**). Clustering coefficients are close to 0, indicating that nodes are quite scattered across the network and not grouped into clusters. The average betweenness is relatively low, suggesting a limited number of central nodes. Additionally, 28% of the network members are connected with at least 3 neighbours, 7% are connected to more than 10 proteins, and 92% of the proteins (N = 1385) are linked to PERK13 through PPI. These data reveal a complex and intricated PPI network, with a consistent part of this network corresponding to genes presenting an ESC1/PERK13-dependent expression (56% of the DEGs). (**Figure 4** and **Table 1**). The organization of the network is composed of modules not related to specific functions, which contrasts with previous observations in studies conducted on other types of biotic interactions, such as plant-pathogen interactions (Delplace et al. 2022; **Figure 4** and **Table S7**). Using the five main classification methods in cytoHubba, we identified hub and bottleneck proteins, with the top ten from each method listed in **Table S11**. Four main hubs were identified in the network of the leaf compartment: MYC2 (AT1G32640), MYB73 (AT4G37260), JAZ1 (AT1G19180), and GI (AT1G22770) (**Figure 4** and **Table S11**). Interestingly, these four genes are mainly known for their role in response to biotic or abiotic stresses, and in jasmonic acid signaling (Fornara *et al*., 2015; Gautam *et al*., 2021; Wang *et al*., 2021; Zhao *et al*., 2024).

**Figure 4.**
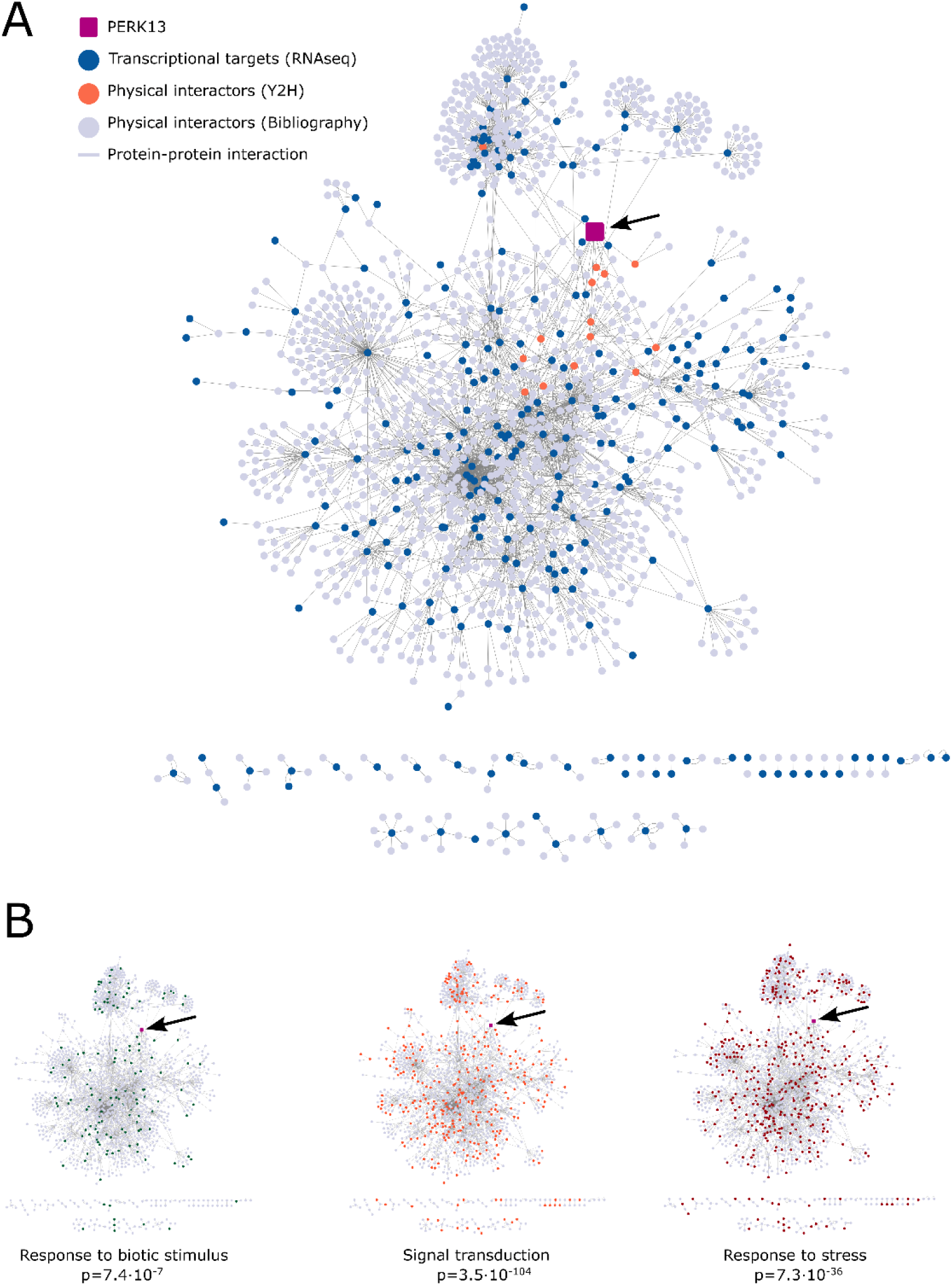
Protein-protein interaction network in leaves is highly interconnected and composed of proteins related to immunity pathways. (A) PERK13 protein-protein interaction networks plotted with Cytoscape showing components used to generate it: PERK13 (magenta, black arrow), PERK13 physical Y2H partners (orange), proteins identified in the RNAseq analysis (blue) and experimental interactors of DEGs and Y2H proteins (grey). (B) Enriched main functional classes in this network showing in color proteins assigned to said category and *p*-value associated.

**Table 1.** General network topology measurements for Leaves and Roots ESC1-dependent networks in response to the presence of *P. annua*. All data were calculated using Network Analyzer of Cytoscape.

| Network | Average degree | Average betweenness | Clustering coefficient | Diameter | Average path length | Number of nodes | Number of edges |
| --- | --- | --- | --- | --- | --- | --- | --- |
| Leaves | 4.03 | 0.015 | 0.033 | 13 | 5.24 | 1502 | 3032 |
| Roots | 2.32 | 0.012 | 0.006 | 10 | 3.81 | 599 | 696 |

The root network exhibits similar characteristics to the leaf network, despite its smaller size, comprising 599 nodes and 696 interactions (**Figure S3**, **Table S7** and **S11**). Interestingly, ESC1/PERK13 appears to be more central and the second main hub in the root compartment (**Figure S3 and S4**).

## Discussion

The molecular mechanisms underlying plant competitive response remain poorly understood. Yet, understanding these mechanisms could help to predict the dynamics of natural plant communities in ecological time (Pierik *et al*., 2013; Frachon *et al*., 2017) and to accelerate breeding programs for the development of crop varieties with enhanced tolerance or competitive ability against weeds (Worthington and Reberg-Horton, 2013; Becker *et al*., 2023). In our study, we focused our analysis on the escape strategy of *A. thaliana* in response to the neighboring plant *P. annua*, identified as one of the main species occurring in *A. thaliana* natural populations (Frachon *et al*., 2017, 2019).

### *ESC1/PERK13,* a gene underlying the natural variation of interspecific competitive response in *A. thaliana*

Previously fine-mapped in a GWAS, a QTL associated with natural variation in the escape strategy, served as the starting point for identifying genes related to plant response to the presence of a competitor. By combining genetic and molecular approaches, we functionally validated the gene underlying this QTL, which encodes a proline-rich extensin-like receptor kinase (PERK), *ESC1*/*PERK13.* To our knowledge, this is the first gene underlying the natural variation of the response to interspecific competition that has been cloned. In the broader context of interspecific plant-plant interactions, only six genes, all conferring resistance to a parasitic plant, have been cloned and functionally validated to date (Li and Timko, 2009; Cardoso *et al*., 2014; Hegenauer *et al*., 2016; Gobena *et al*., 2017; Duriez *et al*., 2019).

Interestingly, *ESC1/PERK13* shares similar eco-genomic patterns with genes involved in plant immunity. For instance, as previously observed for disease resistance genes such as the *NLR* gene *RPS5* and the atypical kinase *RKS1* (Huard-Chauveau *et al*., 2013; Karasov *et al*., 2014; Frachon *et al*., 2017), *ESC1/PERK13* exhibits two highly divergent haplotypes (Haplotype 1 + Haplotype 2 *vs* Haplotype 3) within the French local mapping population TOU-A. The maintenance of this long-lived polymorphism may result from either a fitness cost associated with one of the two haplotypes, conditioned by the presence of competitors, or each haplotype is beneficial in the face of different competitors. In our study, one of the two haplotypes is involved in the response to the presence of *P. annua* and *T. aestivum*. Whether Haplotype 3 is involved in the response to different competitors remains to be tested.

### *ESC1/PERK13* may mediate plant competitive response by perceiving environmental modifications

ESC1/PERK13 is a plasma membrane protein anchored in the cell wall, which is a negative regulator of root hair growth (Won *et al*., 2009; Hwang *et al*., 2016). In the context of plant-plant interactions, this localization suggests that ESC1/PERK13 may trigger an adaptive response by perceiving neighboring plants belowground (either directly or indirectly). Interestingly, *PERK13* expression is upregulated in response to phosphate, iron and nitrogen depletion and *PERK13* modulates root growth under phosphate deficiency (Xue *et al*., 2021). We may therefore conceive that specific nutrient depletion triggered by the proximity of the roots of *P. annua* would provide a feature perceived by ESC1/PERK13 at the root level. This hypothesis would be in line with the common assumption that plants passively perceive their neighbors by detecting the changes in their immediate environment (Novoplansky, 2009). Previous transcriptomic studies indeed revealed the implication of light perception and nutrient-dependent signaling pathways as key components of the plant competition response (Geisler *et al*., 2012; Schmid *et al*., 2013; Horvath *et al*., 2015). However, although light perception and nutrient-related pathways were present in our transcriptomics analysis, biotic stress responses and their associated signaling pathways were identified as the predominant biological processes mediating the response to *P. annua*. Interestingly, over-representation of genes associated with defense responses were also identified in corn in response to weeds (Horvath *et al*., 2018).

The implication of genes related to pathogen response is consistent with the identification of several RLKs in association genetics studies performed across the genome of *A. thaliana* in the context of interspecific interactions (Frachon *et al*., 2019; Libourel *et al*., 2021). These RLKs include Flagellin-Sensing 2 (FLS2), BRI1 Associated Receptor Kinase (BAK1), and BAK1-Interacting Receptor-Like Kinase 1 (BIR1), which belong to distinct subfamilies and participate in different aspects of plant immunity. Their involvement suggests that multiple layers of receptor-mediated signaling, including perception, signal integration, and regulation, could be recruited during plant-plant interactions, supporting the view of a complex and coordinated active plant response.

ESC1/PERK13 belongs to the PERK gene family comprising 15 members in *A. thaliana,* in which it is closely related to the pollen-specific *PERK11* and *PERK12* genes (Nakhamchik *et al*., 2004; Invernizzi *et al*., 2022). While PERKs have traditionally been linked to developmental processes (Borassi *et al*., 2016), emerging evidence indicates a broader role as sensors of abiotic and biotic cues (Invernizzi *et al*., 2022). For instance, while Z15 has been implicated in rice tolerance to low temperatures (Feng *et al*., 2019), the PERK GmHAK2 was identified as a component of the response to herbivores in soybean (Uemura *et al*., 2020), and four *BnPERKs* were downregulated in *Brassica napus* during *Plasmodiophora brassicae* infection (Zhang *et al*., 2025). RLKs interact with one or several co-receptors to fulfill their functions. In this context, the homodimerization of GmHAK2, involved in soybean defense responses to an herbivore, was demonstrated (Bai *et al*., 2009). Still, a screen for interactors using the PERK extracellular domain has not been reported yet. Moreover, PERKs mode of action, notably the identification of their ligands, remains elusive, although they have been hypothesized to be associated with cell wall compounds because of their extracellular proline-rich domain (Nakhamchik *et al*., 2004; Humphrey *et al*., 2015). It was shown that some secondary metabolites in root exudates in soil can depolarize root cell membranes of cucumber seedlings (Maffei *et al*., 2001). We might hypothesize that non-volatile root exudate molecules produced by *P. annua* and *T. aestivum* could depolarize root cell membranes, which in turn would be detected by transmembrane proteins such as PERKs. Identification of these exudate molecules, that induces the escape strategy in *A. thaliana*, would therefore help for a better understanding of the *PERK13* functions.

### ESC1/PERK13 mediates competitive response by distinct root and leaf transcriptomic responses

Based on our transcriptomic analysis, the presence of *P. annua* appears to trigger two distinct reprogramming responses in *A. thaliana* between roots and leaves. This is well illustrated by the few numbers of common DEGs as well as the few biological functions enriched in both compartments. However, biotic and abiotic stresses responses are triggered in both leaves and roots. Interestingly, the ESC1/PERK13-mediated transcriptional changes are more pronounced in leaves than in roots, aligning with the observed aboveground phenotype (i.e., the HD ratio), despite *ESC1*/*PERK13* expression having been previously reported primarily in the root compartment (Hwang *et al*., 2016). The smaller number of genes differentially expressed in roots, mostly linked to perception and transduction, could suggest a key role of this compartment in the perception of *P. annua* and the transfer of a putative signal in the shoot compartment. Accordingly, the overexpression of *ESC1/PERK13* in roots (PERK13Ox) induces an important transcriptomic response in roots, both in the absence and presence of *P. annua*, but also in leaves with 1578 down-regulated and 1165 up-regulated genes specifically in the presence of *P. annua* (Figure S2). This mobile signal could trigger genetic reprogramming in the leaves, thereby leading to a reallocation of resources to stem growth and flowering (Tsikou *et al*., 2018; Lebedeva *et al*., 2020; Dong *et al*., 2022; Zhang *et al*.). In *Populus euphratica*, a similar expression pattern has been observed in response to drought, where root and leaf responses were distinct, with major transcriptional reprogramming occurring in the leaves to regulate photosynthesis (Jiao *et al*., 2021). Interestingly, the limited overlap between responses observed in roots and leaves was also observed during intraspecific competition in *A. thaliana* (Masclaux *et al*., 2012). In this study, photosynthesis-related genes were regulated in the leaves, while nutrient-related genes were regulated in the roots. However, it should be underlined here that biotic stress and defense pathways were activated in both compartments, although the specific genes involved differed. Collectively, these observations suggest a perception of the presence of a neighboring species at the root level, followed by an above-ground transcriptional reprogramming, both processes implicating some functions related to response to biotic stresses. From this perspective, one could hypothesize that signaling compounds from *P. annua* (and *T. aestivum*) root exudates are perceived through ESC1/PERK13, that is mainly expressed in roots. Indeed, such a level of specificity was already observed at intra- or interspecific level (Semchenko *et al*., 2014; Yang *et al*., 2018). Volatile organic compounds (VOCs) are also candidates for the ESC1/PERK13-mediaed response to *P. annua* as they play important roles in plant-plant interactions, even VOCs produced at the root level (Brosset and Blande, 2022). Further experiments compartimenting root and shoot systems are required to determine whether the interactions with *A. thaliana* occur below- or aboveground, as well as the nature of the signaling compound.

### ESC1/PERK13 as an entry node to induce a plant competitive response

Years of global studies have produced extensive data, requiring integration and analysis to better understand the mechanisms involved (Becker *et al*., 2023). Thus, systems biology approaches have been employed to gain deeper insight into the regulation of processes involved in the response to environmental cues, such as pathogen attacks, ultimately leading to the identification of novel immune pathways (Tsuda and Katagiri, 2010; Brauer *et al*., 2018; Delplace *et al*., 2022). Identification of DEGs and ESC1/PERK13 interactors by Y2H screening allowed us to generate two protein-protein interaction networks highlighting the ESC1/PERK13-dependent responses to *P. annua* in the leaf and root compartments. Consistent with our transcriptomics analysis, the two networks displayed a substantial difference in the number of proteins. Nonetheless, in both networks, we identified overrepresented functions, such as signal transduction, shoot system development, and cell communication. These functions are often organized into highly connected modules in other biotic stress-generated networks (Delplace *et al*., 2020). For example, in the case of PTI, four major hormonal sectors were identified which, through compensatory mechanisms under pathogenic perturbations, resulted in network robustness (Kim *et al*., 2014; Hillmer *et al*., 2017). In our study, the nodes of the networks are more dispersed than clustered. The presence of a scale-free topology, combined with functional redundancy, likely contributes to network stability under environmental perturbations (Rodrigues, 2025).

Still, both networks possess major hubs that may be important for network function (Vandereyken *et al*., 2018). For instance, TCP14 is highly connected to many transcription factors and has been identified in pathogen susceptibility networks (Mukhtar *et al*., 2011; Weßling *et al*., 2014). ESC1/PERK13 appears as a bottleneck protein in our study, suggesting it might connect different nodes and modules, possibly influencing the flow of the network. As a receptor potentially acting as a bridge between external stimuli and intracellular responses, ESC1 might indeed have a crucial position in the network. Such a pattern has previously been shown for other receptor proteins (Ahmed *et al*., 2018). However, ESC1/PERK13 interacts only with a few proteins in the network proteins including fifteen Y2H putative interactors identified in our study, and one detected in another analysis, KIPK1 (Humphrey *et al*., 2015), in comparison, for instance, with the well-known FLS2 or BAK1 receptors that have many interactors (Ahmed *et al*., 2018). This could be explained by the lack of data available on plant-plant interactions in comparison with other types of biotic interactions, thereby limiting the network complexity. It could also be possible that the main interactors of ESC1 belong to upstream events of genetic reprogramming. This reinforces the importance of combining methods for identifying signaling pathways and the importance of finding interacting partners of proteins.

## Conclusion

Our work provides the first example of the identification and validation of a gene involved in the natural variation of plant response to competition by a weed species. While plant-plant interactions are thought to be mainly driven by competition for light, nutrients, and water (Pierik *et al*., 2013), we demonstrated in this study that *A. thaliana* could directly or indirectly sense and respond to specific neighborhoods through receptor-ligand proteins, as previously shown in plant-parasitic plant interactions (Li and Timko, 2009; Hegenauer *et al*., 2016; Duriez *et al*., 2019). In addition, we found that many genes associated with plant defense and stress responses were mobilized as part of the escape strategy, reinforcing the concept of an “active” perception and response to a plant competitor.

## Supplementary data

Figure S1. Transcriptionnal reprogramming in the overexpressing line across time and in response to *P. annua*.

Figure S2. *PERK13Ox* DEGs are affected by the presence of *P. annua* only in leaves.

Figure S3. Protein-protein interaction networks in roots is smaller but similar to shoot in terms of classes.

Figure S4. Biological functions are scattered along the two protein-protein interaction networks.

Table S1. List of T-DNA mutant lines used to identify the gene underlying the QTL.

Table S2. Oligonucleotides used in this study.

Table S3. Species used in competition with *A. thaliana*.

Table S4. Characteristics of the experiments performed in this study.

Table S5. Variation among Col-0, perk13.1 and PERK13Ox lines for the response of HD ratio to the presence of seven neighboring species.

Table S6. Lists of genes differentially expressed in leaves and roots 21 days after sowing.

Table S7. Data used to build the root and leaf networks.

Table S8. Treatment effect (absence vs presence of P. annua) for each line.

Table S9. Treatment effect (absence vs presence of P. annua) for the Col-0, perk13.1 and complemented lines and two natural accessions (TOU-A1-124 and TOU-A6-61).

Table S10. List of proteins identified by Y2H as putative interactors of PERK13 kinase domain.

Table S11. List of top hub proteins. Rank is attributed by cytoHubba based on degree, betweenness centrality, bottleneck, closeness centrality and stress measures of the nodes in the leaves and roots networks.

Supplementary Dataset 1: Phenotypic measurements and statistical analyses used in this study

## Acknowledgements

We are grateful to the staff of the LIPME greenhouse for their assistance during the growth chamber experiments. We thank Béatrice Gabinaud for technical help and Sébastien Carrère for assisting with RNAseq analysis.

## Author contributions

F.R., M.H. and D.R. supervised the project. F.R., M.H., D.R., and C.L. designed the experiments. C.L. and F.R. conducted the phenotyping experiments. C.L. analyzed the phenotypic traits. M.I. conducted and analyzed the Y2H screenings. M.H. prepared the samples for RNAseq. M.I. performed the RNAseq analysis and the network recontruction. C.L., M.I., M.H., F.R., and D.R. wrote the manuscript.

## Data Availability

All data supporting the findings of this study are available within the paper and within its supplementary materials published online. RNA-seq reads have been deposited in the NCBI Sequence Read Archive (SRA) under BioProject SRP569129.

## References

1. Ahmed H, Howton TC, Sun Y, Weinberger N, Belkhadir Y, Mukhtar MS. 2018. Network biology discovers pathogen contact points in host protein-protein interactomes. Nature Communications 9, 2312.

2. Asif M, Yang R-C, Navabi A, Iqbal M, Kamran A, Lara EP, Randhawa H, Pozniak C, Spaner D. 2015. Mapping QTL, Selection Differentials, and the Effect of Rht-B1 under Organic and Conventionally Managed Systems in the Attila × CDC Go Spring Wheat Mapping Population. Crop Science 55, 1129–1142.

3. Bai L, Zhang G, Zhou Y, Zhang Z, Wang W, Du Y, Wu Z, Song C-P. 2009. Plasma membrane-associated proline-rich extensin-like receptor kinase 4, a novel regulator of Ca2+ signalling, is required for abscisic acid responses in Arabidopsis thaliana. The Plant Journal 60, 314–327.

4. Baron E, Richirt J, Villoutreix R, Amsellem L, Roux F. 2015. The genetics of intra- and interspecific competitive response and effect in a local population of an annual plant species. (A Bennett, Ed.). Functional Ecology 29, 1361–1370.

5. Becker C, Berthomé R, Delavault P, et al. 2023. The ecologically relevant genetics of plant–plant interactions. Trends in Plant Science 28, 31–42.

6. Borassi C, Sede AR, Mecchia MA, Salgado Salter JD, Marzol E, Muschietti JP, Estevez JM. 2016. An update on cell surface proteins containing extensin-motifs. Journal of Experimental Botany 67, 477–487.

7. Brauer EK, Popescu GV, Singh DK, Calviño M, Gupta K, Gupta B, Chakravarthy S, Popescu SC. 2018. Integrative network-centric approach reveals signaling pathways associated with plant resistance and susceptibility to Pseudomonas syringae. PLOS Biology 16, e2005956.

8. Brosset A, Blande JD. 2022. Volatile-mediated plant–plant interactions: volatile organic compounds as modulators of receiver plant defence, growth, and reproduction. Journal of Experimental Botany 73, 511–528.

9. Cardoso C, Zhang Y, Jamil M, et al. 2014. Natural variation of rice strigolactone biosynthesis is associated with the deletion of two MAX1 orthologs. Proceedings of the National Academy of Sciences 111, 2379–2384.

10. Chin C-H, Chen S-H, Wu H-H, Ho C-W, Ko M-T, Lin C-Y. 2014. cytoHubba: identifying hub objects and sub-networks from complex interactome. BMC Systems Biology 8, S11.

11. Cho H-T, Cosgrove DJ. 2002. Regulation of Root Hair Initiation and Expansin Gene Expression in Arabidopsis. The Plant Cell 14, 3237–3253.

12. Clough SJ, Bent AF. 1998. Floral dip: a simplified method for Agrobacterium -mediated transformation of Arabidopsis thaliana. The Plant Journal 16, 735–743.

13. Coleman RK, Gill GS, Rebetzke GJ. 2001. Identification of quantitative trait loci for traits conferring weed competitiveness in wheat (Triticum aestivum L.). Australian Journal of Agricultural Research 52, 1235–1246.

14. Czechowski T, Stitt M, Altmann T, Udvardi MK, Scheible W-R. 2005. Genome-Wide Identification and Testing of Superior Reference Genes for Transcript Normalization in Arabidopsis. Plant Physiology 139, 5–17.

15. Delplace F, Huard-Chauveau C, Berthomé R, Roby D. 2022. Network organization of the plant immune system: from pathogen perception to robust defense induction. The Plant Journal 109, 447–470.

16. Delplace F, Huard-Chauveau C, Dubiella U, Khafif M, Alvarez E, Langin G, Roux F, Peyraud R, Roby D. 2020. Robustness of plant quantitative disease resistance is provided by a decentralized immune network. Proceedings of the National Academy of Sciences 117, 18099–18109.

17. Dong Q, Hu B, Zhang C. 2022. microRNAs and Their Roles in Plant Development. Frontiers in Plant Science 13.

18. Dorn LA, Pyle EH, Schmitt J. 2000. Plasticity to Light Cues and Resources in Arabidopsis Thaliana: Testing for Adaptive Value and Costs. Evolution 54, 1982–1994.

19. Duriez P, Vautrin S, Auriac M-C, et al. 2019. A receptor-like kinase enhances sunflower resistance to Orobanche cumana. Nature Plants 5, 1211–1215.

20. Feng P, Shi J, Zhang T, et al. 2019. Zebra leaf 15, a receptor-like protein kinase involved in moderate low temperature signaling pathway in rice. Rice 12, 83.

21. Fornara F, de Montaigu A, Sánchez-Villarreal A, Takahashi Y, Ver Loren van Themaat E, Huettel B, Davis SJ, Coupland G. 2015. The GI–CDF module of Arabidopsis affects freezing tolerance and growth as well as flowering. The Plant Journal 81, 695–706.

22. Frachon L, Libourel C, Villoutreix R, et al. 2017. Intermediate degrees of synergistic pleiotropy drive adaptive evolution in ecological time. Nature Ecology & Evolution 1, 1551–1561.

23. Frachon L, Mayjonade B, Bartoli C, Hautekèete N-C, Roux F. 2019. Adaptation to Plant Communities across the Genome of Arabidopsis thaliana. (S Wright, Ed.). Molecular Biology and Evolution 36, 1442–1456.

24. Gautam JK, Giri MK, Singh D, Chattopadhyay S, Nandi AK. 2021. MYC2 influences salicylic acid biosynthesis and defense against bacterial pathogens in Arabidopsis thaliana. Physiologia Plantarum 173, 2248–2261.

25. Geisler M, Gibson DJ, Lindsey KJ, Millar K. 2012. Upregulation of photosynthesis genes, and down-regulation of stress defense genes, is the response of Arabidopsis thaliana shoots to intraspecific competition. Botanical Studies 53, 12.

26. Gobena D, Shimels M, Rich PJ, Ruyter-Spira C, Bouwmeester H, Kanuganti S, Mengiste T, Ejeta G. 2017. Mutation in sorghum LOW GERMINATION STIMULANT 1 alters strigolactones and causes Striga resistance. Proceedings of the National Academy of Sciences 114, 4471–4476.

27. Goldberg DE, Barton AM. 1992. Patterns and Consequences of Interspecific Competition in Natural Communities: A Review of Field Experiments with Plants. The American Naturalist 139, 771–801.

28. Hegenauer V, Fürst U, Kaiser B, Smoker M, Zipfel C, Felix G, Stahl M, Albert M. 2016. Detection of the plant parasite Cuscuta reflexa by a tomato cell surface receptor. Science 353, 478–481.

29. Hillmer RA, Tsuda K, Rallapalli G, Asai S, Truman W, Papke MD, Sakakibara H, Jones JDG, Myers CL, Katagiri F. 2017. The highly buffered Arabidopsis immune signaling network conceals the functions of its components. PLOS Genetics 13, e1006639.

30. Horvath DP, Bruggeman S, Moriles-Miller J, Anderson JV, Dogramaci M, Scheffler BE, Hernandez AG, Foley ME, Clay S. 2018. Weed presence altered biotic stress and light signaling in maize even when weeds were removed early in the critical weed-free period. Plant Direct 2, e00057.

31. Horvath DP, Hansen SA, Moriles-Miller JP, Pierik R, Yan C, Clay DE, Scheffler B, Clay SA. 2015. RNAseq reveals weed-induced PIF3-like as a candidate target to manipulate weed stress response in soybean. New Phytologist 207, 196–210.

32. Huard-Chauveau C, Perchepied L, Debieu M, Rivas S, Kroj T, Kars I, Bergelson J, Roux F, Roby D. 2013. An Atypical Kinase under Balancing Selection Confers Broad-Spectrum Disease Resistance in Arabidopsis. PLOS Genetics 9, e1003766.

33. Humphrey TV, Haasen KE, Aldea-Brydges MG, Sun H, Zayed Y, Indriolo E, Goring DR. 2015. PERK–KIPK–KCBP signalling negatively regulates root growth in Arabidopsis thaliana. Journal of Experimental Botany 66, 71–83.

34. Hwang Y, Lee H, Lee Y-S, Cho H-T. 2016. Cell wall-associated ROOT HAIR SPECIFIC 10, a proline-rich receptor-like kinase, is a negative modulator of Arabidopsis root hair growth. Journal of Experimental Botany 67, 2007–2022.

35. Invernizzi M, Hanemian M, Keller J, Libourel C, Roby D. 2022. PERKing up our understanding of the proline-rich extensin-like receptor kinases, a forgotten plant receptor kinase family. New Phytologist 235, 875–884.

36. Jiao P, Wu Z, Wang X, Jiang Z, Wang Y, Liu H, Qin R, Li Z. 2021. Short-term transcriptomic responses of Populus euphratica roots and leaves to drought stress. Journal of Forestry Research 32, 841–853.

37. Karasov TL, Kniskern JM, Gao L, et al. 2014. The long-term maintenance of a resistance polymorphism through diffuse interactions. Nature 512, 436–440.

38. Kim Y, Tsuda K, Igarashi D, Hillmer RA, Sakakibara H, Myers CL, Katagiri F. 2014. Mechanisms Underlying Robustness and Tunability in a Plant Immune Signaling Network. Cell Host & Microbe 15, 84–94.

39. Lebedeva MA, Yashenkova YaS, Dodueva IE, Lutova LA. 2020. Molecular Dialog between Root and Shoot via Regulatory Peptides and Its Role in Systemic Control of Plant Development. Russian Journal of Plant Physiology 67, 985–1002.

40. Li C, He X, Zhu S, et al. 2009. Crop Diversity for Yield Increase. (DQ Fuller, Ed.). PLoS ONE 4, e8049.

41. Li J, Timko MP. 2009. Gene-for-Gene Resistance in Striga-Cowpea Associations. Science 325, 1094–1094.

42. Libourel C, Baron E, Lenglet J, Amsellem L, Roby D, Roux F. 2021. The Genomic Architecture of Competitive Response of Arabidopsis thaliana Is Highly Flexible Among Plurispecific Neighborhoods. Frontiers in Plant Science 12.

43. Lin C-Y, Chin C-H, Wu H-H, Chen S-H, Ho C-W, Ko M-T. 2008. Hubba: hub objects analyzer—a framework of interactome hubs identification for network biology. Nucleic Acids Research 36, W438–W443.

44. Maere S, Heymans K, Kuiper M. 2005. BiNGO: a Cytoscape plugin to assess overrepresentation of Gene Ontology categories in Biological Networks. Bioinformatics 21, 3448–3449.

45. Maffei M, Camusso W, Sacco S. 2001. Effect of *Mentha* × *piperita* essential oil and monoterpenes on cucumber root membrane potential. Phytochemistry 58, 703–707.

46. Martorell C, Freckleton RP. 2014. Testing the roles of competition, facilitation and stochasticity on community structure in a species-rich assemblage. Journal of Ecology 102, 74–85.

47. Masclaux FG, Bruessow F, Schweizer F, Gouhier-Darimont C, Keller L, Reymond P. 2012. Transcriptome analysis of intraspecific competition in Arabidopsis thalianareveals organ-specific signatures related to nutrient acquisition and general stress response pathways. BMC Plant Biology 12, 227.

48. Moncada P, Martínez CP, Borrero J, Chatel M, Gauch Jr H, Guimaraes E, Tohme J, McCouch SR. 2001. Quantitative trait loci for yield and yield components in an Oryza sativa×Oryza rufipogon BC2F2 population evaluated in an upland environment. Theoretical and Applied Genetics 102, 41–52.

49. Mukhtar MS, Carvunis A-R, Dreze M, et al. 2011. Independently Evolved Virulence Effectors Converge onto Hubs in a Plant Immune System Network. Science 333, 596–601.

50. Nakhamchik A, Zhao Z, Provart NJ, Shiu S-H, Keatley SK, Cameron RK, Goring DR. 2004. A Comprehensive Expression Analysis of the Arabidopsis Proline-rich Extensin-like Receptor Kinase Gene Family using Bioinformatic and Experimental Approaches. Plant and Cell Physiology 45, 1875–1881.

51. Neve P, Vila-Aiub M, Roux F. 2009. Evolutionary-thinking in agricultural weed management. New Phytologist 184, 783–793.

52. Novoplansky A. 2009. Picking battles wisely: plant behaviour under competition. Plant, Cell & Environment 32, 726–741.

53. Oerke E-C. 2006. Crop losses to pests. The Journal of Agricultural Science 144, 31–43.

54. Pierik R, Mommer L, Voesenek LA. 2013. Molecular mechanisms of plant competition: neighbour detection and response strategies. Functional Ecology 27, 841–853.

55. Robinson MD, McCarthy DJ, Smyth GK. 2010. edgeR: a Bioconductor package for differential expression analysis of digital gene expression data. Bioinformatics 26, 139–140.

56. Rodrigues JA. 2025. A Topological Approach to Protein–Protein Interaction Networks: Persistent Homology and Algebraic Connectivity. International Journal of Topology 2, 8.

57. Roux F, Bergelson J. 2016. Chapter Four - The Genetics Underlying Natural Variation in the Biotic Interactions of Arabidopsis thaliana: The Challenges of Linking Evolutionary Genetics and Community Ecology. In: Orgogozo V, ed. Genes and Evolution. Current Topics in Developmental Biology. Academic Press, 111–156.

58. Sato Y, Wuest SE. 2025. The Genetics of Plant–Plant Interactions and Their Cascading Effects on Agroecosystems—from Model Plants to Applications. Plant and Cell Physiology 66, 477–489.

59. Schmid C, Bauer S, Müller B, Bartelheimer M. 2013. Belowground neighbor perception in Arabidopsis thaliana studied by transcriptome analysis: roots of Hieracium pilosella cause biotic stress. Frontiers in Plant Science 4.

60. Semchenko M, Saar S, Lepik A. 2014. Plant root exudates mediate neighbour recognition and trigger complex behavioural changes. New Phytologist 204, 631–637.

61. Subrahmaniam HJ, Libourel C, Journet E-P, Morel J-B, Muños S, Niebel A, Raffaele S, Roux F. 2018. The genetics underlying natural variation of plant-plant interactions, a beloved but forgotten member of the family of biotic interactions. The Plant Journal 93, 747–770.

62. Tilman D. 1985. The Resource-Ratio Hypothesis of Plant Succession. The American Naturalist 125, 827–852.

63. Tsikou D, Yan Z, Holt DB, Abel NB, Reid DE, Madsen LH, Bhasin H, Sexauer M, Stougaard J, Markmann K. 2018. Systemic control of legume susceptibility to rhizobial infection by a mobile microRNA. Science 362, 233–236.

64. Tsuda K, Katagiri F. 2010. Comparing signaling mechanisms engaged in pattern-triggered and effector-triggered immunity. Current Opinion in Plant Biology 13, 459–465.

65. Uemura T, Hachisu M, Desaki Y, et al. 2020. Soy and Arabidopsis receptor-like kinases respond to polysaccharide signals from Spodoptera species and mediate herbivore resistance. Communications Biology 3, 1–11.

66. Van Criekinge W, Beyaert R. 1999. Yeast Two-Hybrid: State of the Art. Biological Procedures Online 2, 1–38.

67. Vandereyken K, Van Leene J, De Coninck B, Cammue BPA. 2018. Hub Protein Controversy: Taking a Closer Look at Plant Stress Response Hubs. Frontiers in Plant Science 9.

68. Walsche AD, Vergne A, Rincent R, Roux F, Nicolas S, Welcker C, Mezmouk S, Charcosset A, Mary-Huard T. 2025. metaGE: Investigating genotype x environment interactions through GWAS meta-analysis. PLOS Genetics 21, e1011553.

69. Wang L, Qiu T, Yue J, et al. 2021. Arabidopsis ADF1 is Regulated by MYB73 and is Involved in Response to Salt Stress Affecting Actin Filament Organization. Plant and Cell Physiology 62, 1387–1395.

70. Weinig C, Johnston J, German ZM, Demink LM. 2006. Local and Global Costs of Adaptive Plasticity to Density in Arabidopsis thaliana. The American Naturalist 167, 826–836.

71. Wender NJ, Polisetty CR, Donohue K. 2005. Density-dependent processes influencing the evolutionary dynamics of dispersal: a functional analysis of seed dispersal in Arabidopsis thaliana (Brassicaceae). American Journal of Botany 92, 960–971.

72. Weßling R, Epple P, Altmann S, et al. 2014. Convergent targeting of a common host protein-network by pathogen effectors from three kingdoms of life. Cell host & microbe 16, 364–375.

73. Won S-K, Lee Y-J, Lee H-Y, Heo Y-K, Cho M, Cho H-T. 2009. cis-Element- and Transcriptome-Based Screening of Root Hair-Specific Genes and Their Functional Characterization in Arabidopsis. Plant Physiology 150, 1459–1473.

74. Worthington M, Reberg-Horton C. 2013. Breeding Cereal Crops for Enhanced Weed Suppression: Optimizing Allelopathy and Competitive Ability. Journal of Chemical Ecology 39, 213–231.

75. Xue C, Li W, Shen R, Lan P. 2021. PERK13 modulates phosphate deficiency-induced root hair elongation in Arabidopsis. Plant Science 312, 111060.

76. Yang X-F, Li L-L, Xu Y, Kong C-H. 2018. Kin recognition in rice (Oryza sativa) lines. New Phytologist 220, 567–578.

77. Zhang Z, Fu T, Zhou C, Liu F, Zeng L, Ren L, Tong C, Liu L, Xu L. 2025. Genome-Wide Analysis of the PERK Gene Family in Brassica napus L. and Their Potential Roles in Clubroot Disease. International Journal of Molecular Sciences 26, 2685.

78. Zhang J, Li Y, Bao Q, Wang H, Hou S. Plant elicitor peptide 1 fortifies root cell walls and triggers a systemic root-to-shoot immune signaling in Arabidopsis. Plant Signaling & Behavior 17, 2034270.

79. Zhao X, He Y, Liu Y, Wang Z, Zhao J. 2024. JAZ proteins: Key regulators of plant growth and stress response. The Crop Journal 12, 1505–1516.

