## Supplementary figures for "An *Arabidopsis* receptor-like kinase mediates competitive plant-plant interactions"

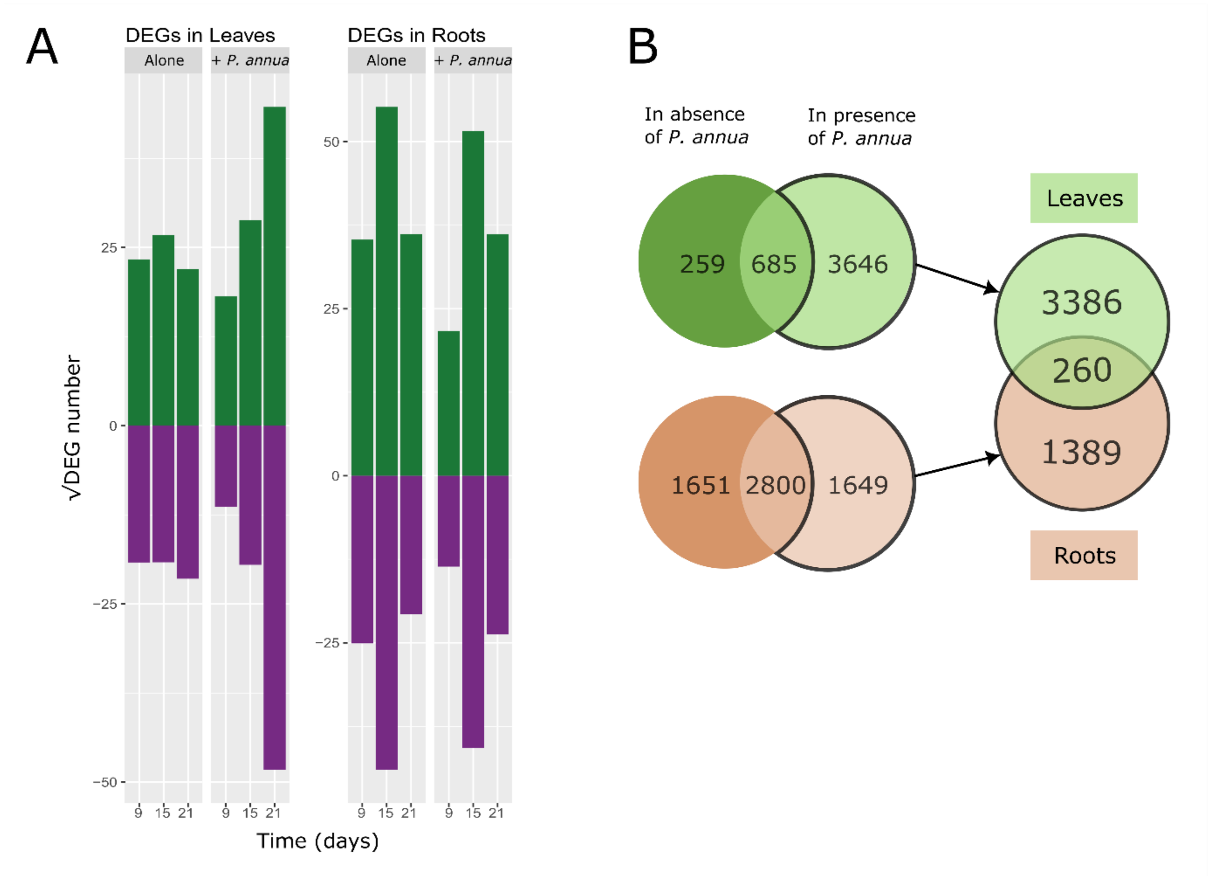


**Figure S1. Transcriptionnal reprogramming in the overexpressing line across time and in response to *P. annua*.** (A) Upregulated genes (green) and downregulated genes (purple) found in the *PERK13*Ox in comparison to Col-0 for each treatment, tissue and timepoint. (B) Venn diagrams representing the DEGs found in21-day-old plants in the absence (dark color) and in the presence of *P. annua* (light color), for leaves (green) and roots (orange). The parts of diagram outlined in bold represent the DEGs specific to *P. annua* response and are used for DEGs comparison between leaves and roots.

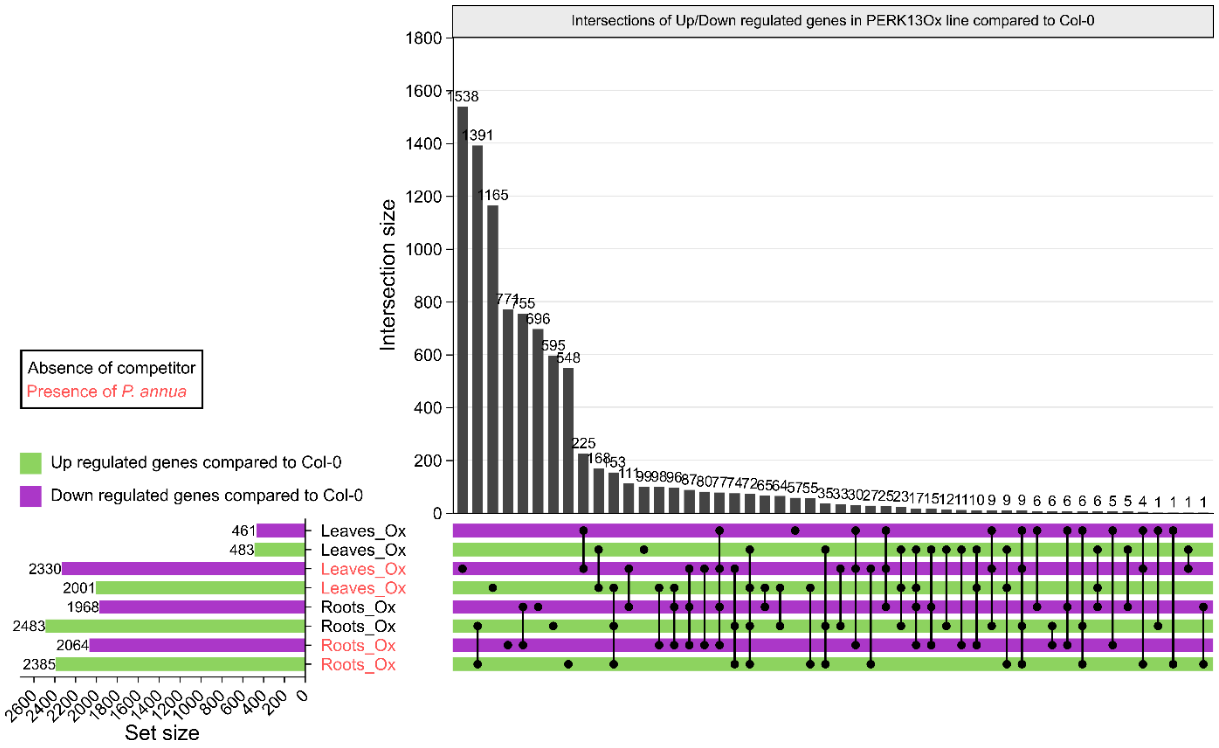


**Figure S2. *PERK13Ox* DEGs are affected by the presence of *P. annua* only in leaves.** Upset plot of up (green) and down (purple) regulated DEGs of *PERK13*Ox compared to Col-0 identified at 21das in absence (black) or presence (red) of *P. annua*.

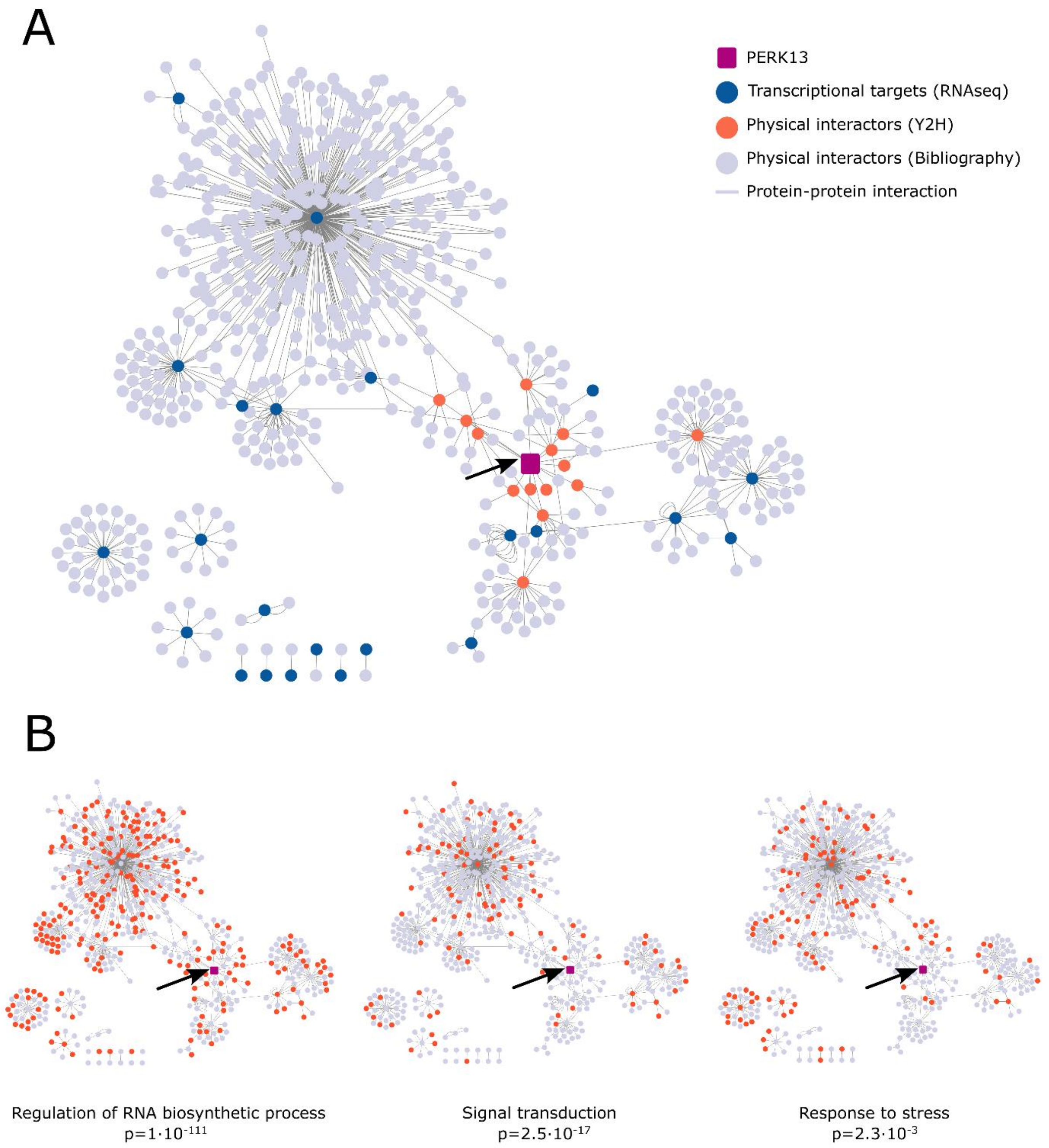


**Figure S3. Protein-protein interaction networks in roots is smaller but similar to shoot in terms of classes.** (A) PERK13 protein-protein interaction networks plotted with Cytoscape showing components used to generate it: PERK13 (magenta, black arrow), PERK13 physical Y2H partners (orange), proteins identified in the RNAseq analysis (blue) and experimental interactors of DEGs and Y2H proteins (grey). (B) Enriched main functional classes in this network showing in color proteins assigned to said categories.

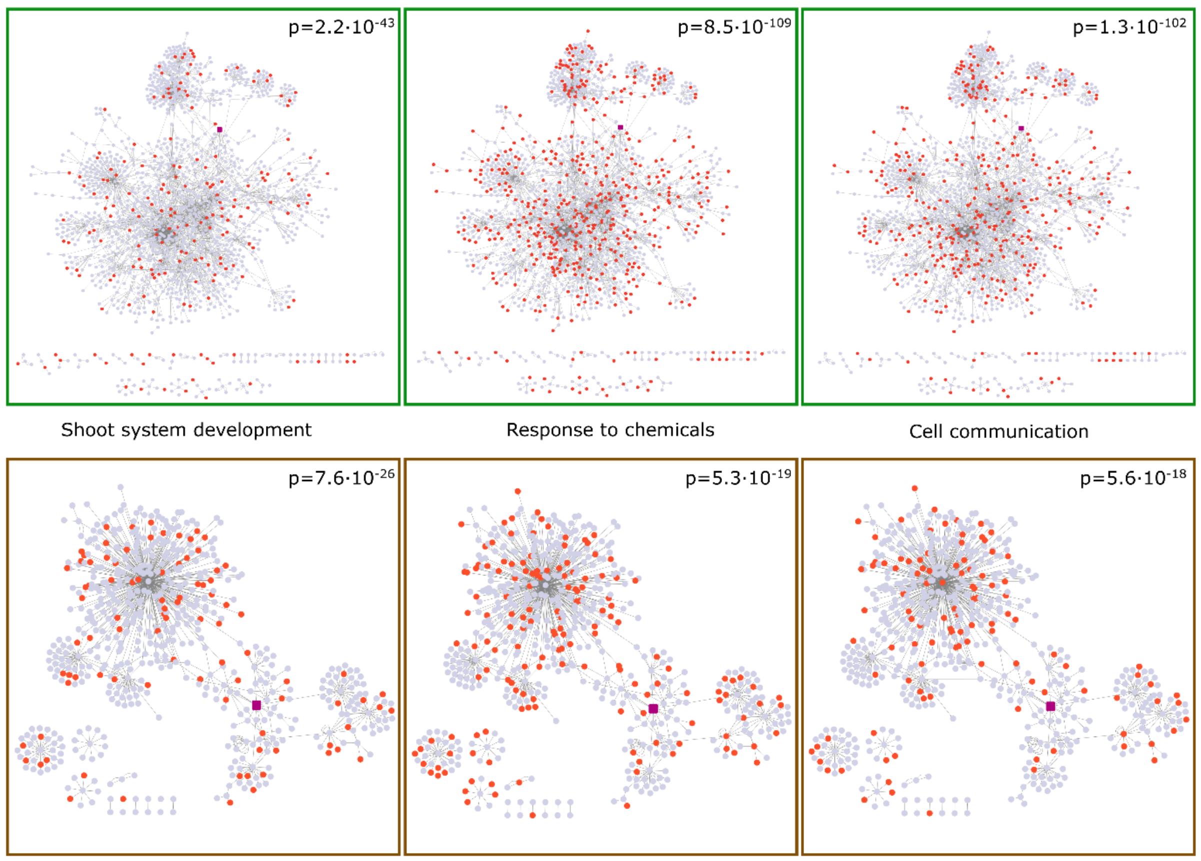


**Figure S4. Biological functions are scattered along the two protein-protein interaction networks.** GO terms enriched in leaves (green) or roots (brown) networks and associated p-values. Nodes associated with the categories are highlighted in orange.
